# Mapping tree species for restoration potential resilient to climate change

**DOI:** 10.1101/2021.06.04.447113

**Authors:** Nina van Tiel, Lisha Lyu, Fabian Fopp, Philipp Brun, Johan van den Hoogen, Dirk Nikolaus Karger, Niklaus E. Zimmermann, Thomas W. Crowther, Loïc Pellissier

## Abstract

The restoration of forest ecosystems is associated with key benefits for biodiversity and ecosystem services. Where possible, ecosystem restoration efforts should be guided by a detailed knowledge of the native flora to regenerate ecosystems in a way that benefits natural biodiversity, ecosystem services, and nature’s contribution to people. Machine learning can map the ecological suitability of tree species globally, which then can guide restoration efforts, especially in regions where knowledge about the native tree flora is still insufficient. We developed an algorithm that combines ecological niche modelling and geographic distributions that allows for the high resolution (1km) global mapping of the native range and suitability of 3,987 tree species under current and future climatic conditions. We show that in most regions where forest cover could be potentially increased, heterogeneity in ecological conditions and narrow species niche width limit species occupancy, so that in several areas with reforestation potential, a large amount of potentially suitable species would be required for successful reforestation. Local tree planting efforts should consider a wide variety of species to ensure that the equally large variety of ecological conditions can be covered. Under climate change, a large fraction of the surface for restoration will suffer significant turnover in suitability, so that areas that are suitable for many species under current conditions will not be suitable in the future anymore. Such a turnover due to shifting climate is less pronounced in regions containing species with broader geographical distributions. This indicates that if restoration decisions are solely based on current climatic conditions, a large fraction of the restored area will become unsuitable in the future. Decisions on forest restoration should therefore take the niche width of a tree species into account to mitigate the risk of climate-driven ecosystem degradation.

## Introduction

The restoration of natural ecosystems benefits biodiversity and ecosystem services (Groot et al. 2013; R. Chazdon and Brancalion 2019). The UN Decade on Ecosystem restoration has begun to catalyze interest in nature restoration, with the goal of addressing biodiversity loss and climate change, and at the same time enhancing human wellbeing across the globe (Fischer et al. 2021). A key component in this global effort is the restoration of forests, which represent the largest repositories of biodiversity and carbon on land (Lewis et al. 2019). In many cases, the most effective approaches for forest restoration involve the protection of land so that natural forests can recover naturally (Parrotta, Turnbull, and Jones 1997; Crouzeilles et al. 2017). Many tree species are associated with essential ecosystem services including carbon sequestration, and climate regulation (Pan et al. 2011) and offering habitats for many other organisms (Zellweger et al. 2013). In many other cases, active restoration practices such as soil amendment, tree planting, applied nucleation and agroforestry practices can be valuable for facilitating the speed at which biodiversity can recover (Zahawi et al. 2013) and could provide an efficient means to improve biodiversity globally (Kemppinen et al. 2020). To avoid the many potential pitfalls of planting trees, including planting problematic (e.g. potentially invasive) tree species, restoration efforts should be guided by solid information on species’ native distribution range, ecological preference and resistance to climate change (MacKenzie and Mahony 2021). Indeed, with the high velocity of climate change, deciding on which tree species to use for assisted restoration corresponds to aiming at a moving target, where planting under suitable conditions for a species, does not guarantee that they will persist as the tree grows (Gray and Hamann 2011). Hence, climate-smart ecosystem restoration requires a fundamental understanding of species’ native ranges as well as species ecological suitabilities under both current and future environmental conditions.

Active restoration initiatives face considerable social and ecological challenges that must be overcome in order to generate sustainable ecosystems (Toledo et al. 2018; Holl 2017). Among these, a top priority is the identification of tree species that are native to the region (Lu et al. 2017), to avoid generating reforested systems that are maladapted for coexistence with the local biodiversity (Meli et al. 2014). In addition, the ecological preferences of species selected should fit the local environmental conditions to ensure long-term survival and reproduction and thus a successful restoration of natural ecosystems (Shono, Davies, and Chua 2007). Global ecological studies have focussed primarily on predicting patterns of aggregate community-level metrics such as species diversity (Kreft and Jetz 2007), biomass or carbon stock (Pan et al. 2011). Yet, identifying the species that are suitable to a target region for restoration requires the consideration of species-level environmental tolerances and species ranges across thousands of independent taxa. Given that our Earth is home to over 60,000 tree species (Beech et al. 2017), understanding the spatial distribution of each remains a considerable research challenge, but one that urgently needs to be addressed to guide ecologically responsible active ecosystem reforestation efforts. To overcome this challenge, machine learning can support the mapping of species habitat suitability (Pecchi et al. 2019). In particular, species distribution modelling (Guisan and Zimmermann 2000) can inform ecosystem management decisions and support biodiversity conservation (Guisan et al. 2013). However, until now, the studies exploring the distributions of tree species at high spatial resolutions has been limited to individual tree species and local to regional scales (Hidalgo et al. 2008; Vessella and Schirone 2013). Modeling the distribution of any individual species globally requires a massive effort to combine datasets on occurrences (and preferably absences) from a wide variety of sources. Moreover, doing so for thousands of species represents an unprecedented challenge in handling large datasets, and computing model fits and global distribution maps with a resolution that is high enough for meaningful reforestation decisions.

The challenge of tree species selection for restoration is additionally compounded by changing climatic conditions (Gray and Hamann 2011). A decision that seems optimal under current climatic conditions, might actually not be the best management solution for the future due to the rapid shift in climatic conditions (Ravenscroft et al. 2010). The resulting shifts in potential plant distributions mean that species currently thriving in a location may not necessarily be able to survive there in the coming decades (McKenney et al. 2007; Dyderski et al. 2018) This will lead to shift in species potential composition (Morin et al. 2018) and a possible degradation of the potential ecosystem services provided (Mina et al. 2017). Hence, climate-smart forest and land management necessitates a basic understanding of which species also have the potential to survive under future climate conditions (Charney et al. 2016). Yet, until now, the potential distribution of tree species under current and future climate scenarios remains a major uncertainty in our efforts to manage and restore land effectively, threatening the value of forested land if not sustainably planned (Hanewinkel et al. 2013). In order to inform restoration projects and plan robust replanting solutions, we need forecasts of future suitabilities for tree species. Machine learning approaches not only allow mapping the present distributions of thousands of tree species across the global forest system (Dyderski et al. 2018), but also projecting their response to global changes, when coupled with climate change projections (Karger et al. 2017; Dyderski et al. 2018).

Here, we investigated the native distribution range and ecological suitability of thousands of tree species under present and future climates to inform ecologically responsible ecosystem restoration initiatives. Using a cloud implementation of an environmental niche model algorithms, we generated species distribution models for 3,987 tree species across the globe. We used a global network of forest plots, including thousands of direct species observations, to determine the most frequent and locally abundant tree species in all major bioregions. Using observations from 13 databases (Table 1), we mapped the distribution of 3,987 tree species. We first evaluated how many tree species are necessary to cover all the abiotic conditions in 39 regions of forest restoration potential (Bastin et al. 2019). We further evaluated whether climate change would cause an important loss in suitability for species, which would challenge restoration decisions based solely on current environmental conditions. Specifically, we asked the following questions:

1. How many species are needed to cover a significant fraction of those areas of high reforestation potential given potential local heterogeneity and limited ecological niche breadth of species?
2. How do suitable areas for species change with ongoing climate change, and does it impact the selection of species for restoration presently? ?

**Table 1.** Online databases used as data sources for occurrence data.

| Database name | URL |
| --- | --- |
| Botanical Information and Ecology Network (BIEN) | https://bien.nceas.ucsb.edu/ |
| BIOMASS | https://www.nature.com/articles/sdata201770#data-records |
| Cardullo et al. 2017 | https://www.sciencedirect.com/science/article/pii/S2352340917301981?via%3Dihub |
| CONIFER | https://herbaria.plants.ox.ac.uk/bol/conifers |
| DRYFLOR | http://www.dryflor.info/data/datadownload |
| GBIF | https://www.gbif.org/occurrence/download/0032444-200221144449610 |
| GFBI | https://www.gfbinitiative.org/ |
| INDIABIODIVERSITY | https://indiabiodiversity.org/observation/list |
| IDIGBIO | https://www.idigbio.org/portal/search |
| MUSEUM | Field collection records and manually georeferenced herbarium data from Naturalis (L) and Paris (P) natural history museums (unpublished) |
| PNG | http://www.pngplants.org/search.htm |
| PREDICTS | https://data.nhm.ac.uk/dataset/the-2016-release-of-the-predicts-database |
| RAINBIO | https://gdauby.github.io/rainbio/download_page.html |

We expect that our mapping approach can support projects with the aim of restoring forest ecosystems, informing about the fit of species to present ecological conditions but also the species resilience to climate change, and thus indicating whether ecosystem services associated with forests can be provided sustainably.

## Results

### Species occupancy and richness globally and in selected regions

Based on a global inventory of forest plots with information on relative abundance (Blundo et al. 2021), we selected 4,172 species in all main bioregions globally, which we mapped with a combination of geographic polygons and species distribution modelling. We selected the 50 most frequently observed species from each biome (1,987 species) according to available forest plots, and subsequently selected another 2,185 species that co-occur with a species from the first set in at least 50% the forest plots in which it was reported. The dominant species spanned multiple plant families, with the most frequently represented families being Fabaceae (457 species), Myrtaceae (181 species) and Rubiaceae (171 species). We obtained 3,987 species distributions as species lacking enough data in their reported native ranges were not modelled. Moreover, our results generally support the observations of previous species range mapping efforts (Caudullo, Welk, and San-Miguel-Ayanz 2017), whilst also providing a resource for many areas where tree species ranges are not available, including many tropical areas (Figure 1). Our set of selected species is representative of the global diversity patterns, with more species selected in the tropics compared with higher latitudes, and covers the main potentially forested areas globally (Bastin et al. 2019). Due to this selection process, we may be propagating sampling biases as we see few modelled species in some areas, such as Central Asia (Figure 1), which corresponds with the more limited availability of forest plots in those regions. We found that the geographic boundary was generally important in determining the species range since most species showed suitable areas outside of their geographic limit. Our results show the interest of combining both geographic limit and ecological suitability to map the potential of species to serve for reforestation in local restoration sites focusing on native species.

**Figure 1.**
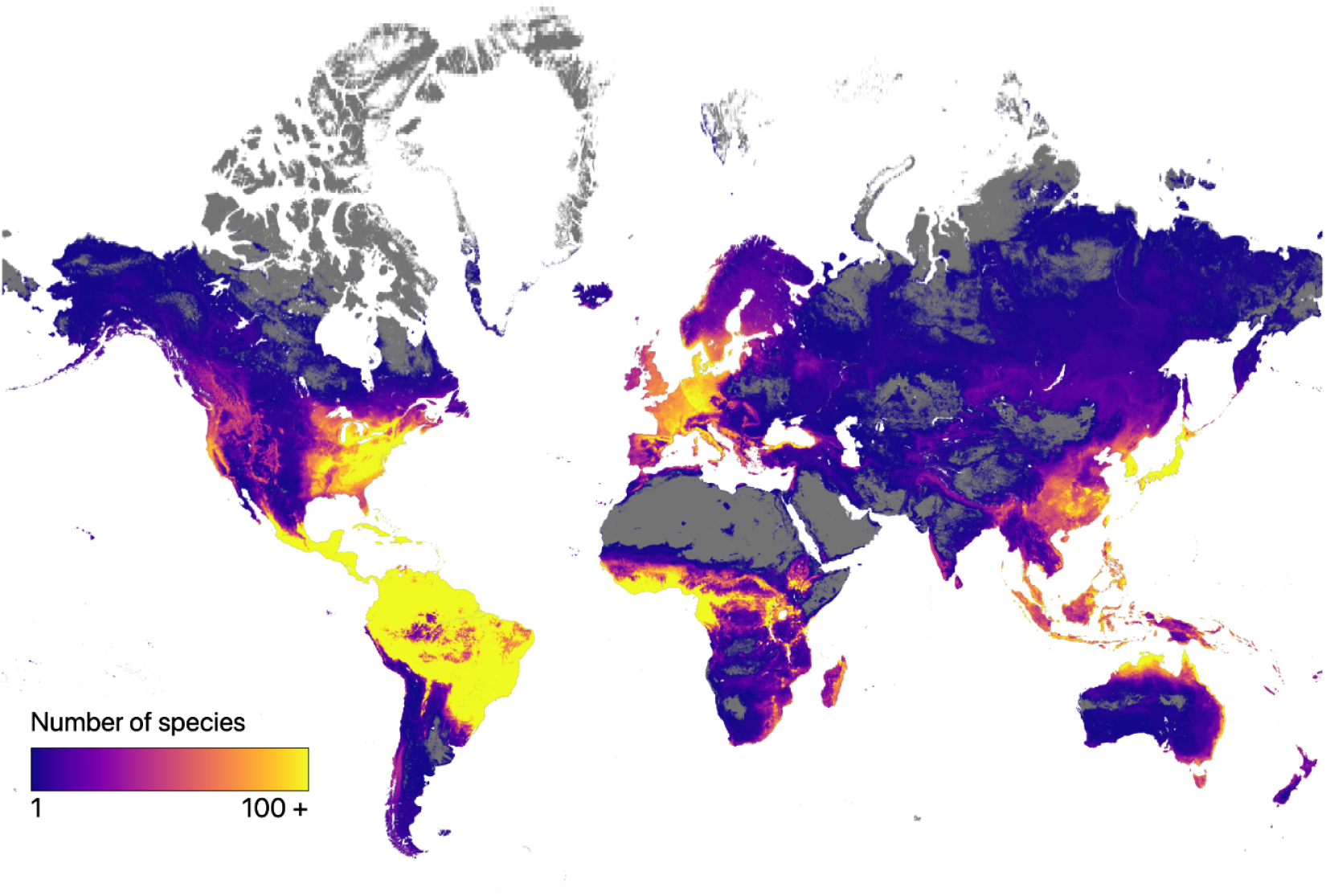
Species richness map obtained by stacking all 3,987 species distribution models. While the richness broadly corresponds with known phytodiversity gradients, the species selection also leads to a bias, including a lower coverage of species on the indian subcontinent, in South East Asia, Russia and an overrepresentation in Europe or North America, where many forest plots are occurring.

While our mapping can address the entire potential area for tree cover, as case studies we specifically selected 39 regions with high reforestation potential, by selecting areas larger than 40,000 km2 that have at least 30% of restorable tree cover (Bastin et al. 2019) and for which our modelled species distributions covered at least 90% of the area. Those regions usually show significant anthropogenic impact and likely a slow natural regeneration. Those regions, when classified by bioregions, have different number of species from our global modelled pool (median number of species per area, Palearctic 82.5, Indomalayan 146.0, Nearctic 167.0, Afrotropic 197.5, Australasia 221.0, Neotropic 1123.0). Moreover, we found that species have generally higher median area coverage in some regions compared with others (median area covered by the distribution of one species, Indomalayan=984 km2, Afrotropic=4’436 km2, Palearctic=4’805 km2, Neotropics=6’145 km2, Australasia=11’363 km2, Nearctic=16’455 km2), suggesting that tree species could have larger environmental niche width in some regions compared with others. Unlike previous large scale modelling efforts (Thuiller et al. 2011), our approach combined both species distribution modelling and the geographic range limits, which can for instance represent dispersal limitations (Merow, Wilson, and Jetz 2017; Polanco et al. 2020). Complementing previous local use of species distribution models to inform forest management (Booth 2018), our modelling synthesis provides a general approach to inform the suitability of potentially dominant and frequent species at local sites and provides structuring species for reforestation in the context of global restoration projects.

### Species cover accumulation curves across selected regions

We found that in most regions a relatively large number of species are needed to allow reforestation of a large fraction of the target areas (Figure 2). Using species accumulation curves, we show that for most regions we need between 2 and 29.6 species to cover half of an area with forest potential. Moreover, the cover accumulation curve showed a saturation in a range between 50 and 200 species (Figure 2), but also indicated a large variation among the regions. We found difference among regions in the number of species needed to reach 95% coverage among species (median number of species required to cover 95% of area, Palearctic=12.75, Nearctic=27.25, Afrotropic=27.75, Neotropic=31.50, Australasia=34.50, Palearctic=12.75, Afrotropic=27.75, Indomalayan=69.00, Australasia=34.50). Some regions saturate faster mainly because they contain species with overall larger potential range size (Figure 2). Indeed, we found a negative correlation between median range size and the number of species needed to cover 95% (Pearson’s r= −0.290). This suggests that regions with species containing large range sizes need a smaller species pool to reach near-complete coverage. Our results suggest that forest restoration generally requires a variety of local trees to cover the diverse environmental conditions contained within a target region. Hence, those maps could serve as a first guide for local stakeholders, which can be combined with local observations and local expert knowledge to finally decide on the tree species to be considered (Gastón et al. 2014).

**Figure 2.**
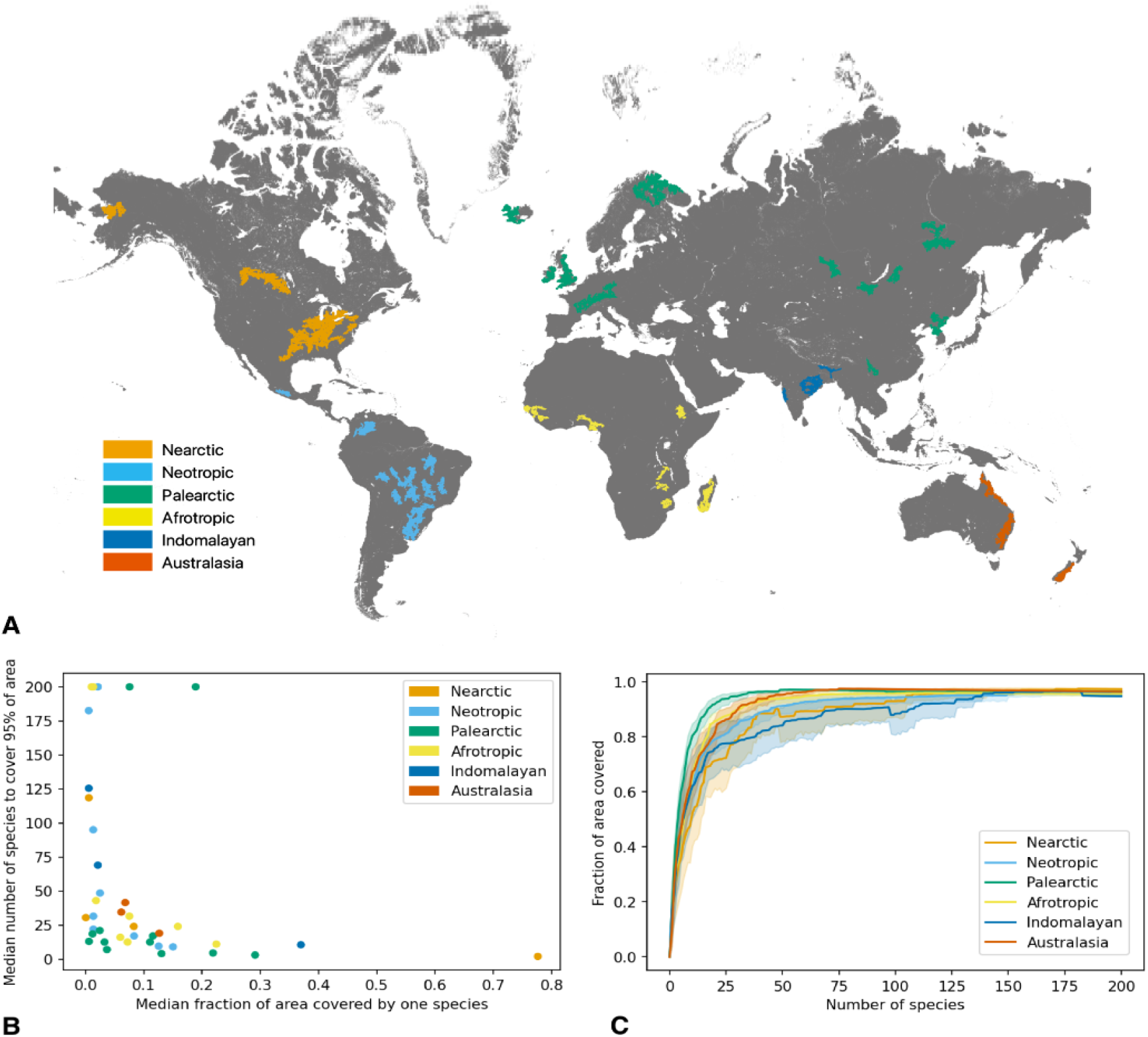
Map of the 39 selected areas as case study to investigate reforestation potential by three species coloured by bioregions (A). Relationship between the median fraction of area covered by a given species and the median number of species to cover 95 percent of the total area (B). Species cover accumulation curve showing the fraction of area covered as the number of species considered increases (C).

### Moving target under climate change

We compared the species distributions mapped for current climate with those projected for predicted future climate for 2041-2070 unser a SSP 5.85 scenario. We aimed to assess the consequence of climate change if reforestation decisions would be solely taken based on present climate. We demonstrate that climate change will cause a shift in species suitability in most regions, challenging the decision on reforestation solely based on present climate (Figure 3). Specifically, we found that climate change caused the decline in median suitable area of species (median proportion of species losing suitable area, Palearctic=0.372, Australasia=0.466, Nearctic=0.529, Afrotropic=0.532, Neotropics=0.638, Indomalayan=0.644) (Figure 3). Despite those declines, the fraction of fully extirpated species from the target area remain relatively low (median proportion of extirpated species, Palearctic=0.021, Australasia=0.032, Afrotropic=0.053, Nearctic=0.092, Neotropics=0.123, Indomalayan=0.164), which indicate mainly a redistribution within the target regions. Moreover, regionally, climate change will not cause a significant decline in areas potentially suitable for any tree species occurring presently, but we nevertheless found significant species turnover at the level of the cell. We found that the proportion of the surface area of the analysed regions for which a least one species will get extirpated is close to one (median proportion of areas where at least one species will disappear under climate change, Indomalayan=0.903, Australasia=0.950, Palearctic=0.957, Nearctic=0.962, Neotropics=0.963, Afrotropic=0.964). Hence, in most areas, a subset of species will show a complete shift from suitable to unsuitable, which will challenge the selection of species for reforestation. Overall, we found a significant turnover of species suitabilities in most cells (median Jaccard index, or similarity of current and future species distributions, Palearctic=0.148, Nearctic=0.191, Neotropics=0.238, Afrotropic=0.333, Indomalayan=0.371, Australasia=0.540). In those areas with high expected impact of climate change, reforestation should take particular care in considering future suitabilities under climate change. We found that different areas are very much unequal in the face of climate change. Our results support empirical evidence, where several regions have already documented species stresses or declines as a consequence of recent climate change (van Wees et al. 2021). We further found a positive correlation between the median potential range size by a single species and the median similarity of the current and future composition of species in a given region (Pearson *r*= 0.470). This suggests that areas with more habitat specialized species of smaller range show stronger compositional turnover under climate change.

**Figure 3.**
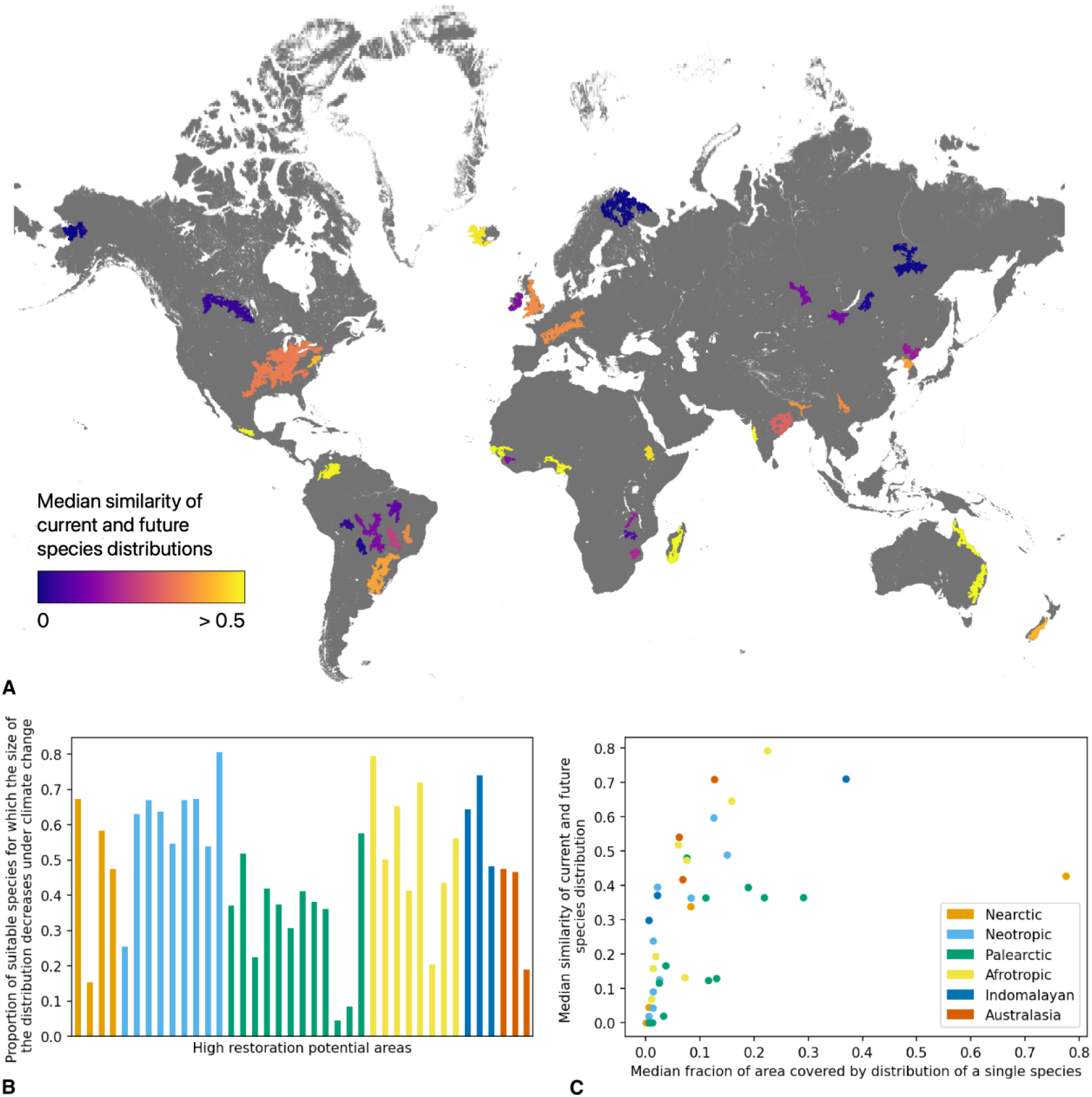
Map of the 39 selected areas coloured by median similarity (Jaccard index) of the current and future distributions of species (A). Proportion of suitable species for which the size of the distribution decreases under climate change for each area (B). Relationship between the median fraction of areas covered by a given species and the median similarity of the current and future species distributions (C).

## Discussion

Given the high structuring importance of tree species on many terrestrial habitats globally, knowledge of tree species distribution is central for restoration. Hence, maximizing the success of active ecosystem restoration and management initiatives across the globe requires fundamental information about which mixtures of species are best suited to surviving in those specific regions (Crouzeilles et al. 2020). By combining species distribution modelling (Guisan and Zimmermann 2000) with spatial polygons to constrain the suitable area to the native range (Polanco et al. 2020), we provide suitability for many tree species, beyond regions where species range maps are already available and broadly used by forester to manage forest resources (Caudullo, Welk, and San-Miguel-Ayanz 2017).

Unlike previous modelling approaches (Rondinini et al. 2011), our pipeline considers both spatial boundaries and ecological suitability and can serve the mapping of organisms beyond tree species (Polanco et al. 2020). Typically, while global species range maps are available for well-known taxonomic groups such as vertebrates (BirdLife International & NatureServe 2015) and from the IUCN (2015) Red list for amphibians and terrestrial mammals), there exist still no high resolution mapping for tree species (Serra-Diaz et al. 2018). By building ~4000 plant species distribution models across the globe, and 39 selected areas as case studies, we demonstrate that it is now possible to map the habitat suitability of tree species at high resolution for restoration purposes.

Passive restoration has been suggested to be a more effective and affordable approach to recover and conserve the ecosystem structure and function (Crouzeilles et al. 2017), but in highly degraded landscapes the rate of natural colonisation could be too slow to regenerate ecosystem over the next couple of decades (Robin L. Chazdon 2008). Hence, selecting native species that fit the local environmental conditions is central to restoration efforts, where active restoration is necessary (Gastón et al. 2014) and could complement local expert knowledge (Suárez et al. 2012). Our approach allowed us to automatically map species ranges. Modelling global distribution ranges for so many species was also allowed by the newly available implementation of machine learning in Google Earth Engine (Gorelick et al. 2017), allowing the coupling of powerful geostatistics and species distribution modelling. As a case study, we selected 39 areas globally and investigated the suitability of species presently and under climate change. We demonstrate that covering the heterogeneous environmental conditions in a region requires a large tree species pool, supporting local assessments for tree nurseries (Vidal et al. 2020). We further found variability in the region in terms of median species cover, which was correlated with the number of species to reach 95% cover of the region. Hence, in regions with species that are geographically more restricted, it requires more species to cover those entire areas. Hence, our results broaden the findings from local studies (Silva et al. 2017; Vidal et al. 2020), showing that forest restoration needs to consider a variety of species regionally, and offer a global information dataset to support future decisions.

Forest restoration projects are not only about restoring the type of forest that used to grow in a region under current climate scenarios, but also about anticipating climate change impact and managing for tree species that are resistant to future environmental change (Prober et al. 2015). In most pixels we found at least one species for which currently suitable conditions are expected to become unsuitable under climate change, implying a high risk of making a wrong decision as regard to restoration. We identified less expected turnover in regions with many high proportion or ranged species with large ranges, indicating that it might be less risky to use broadly distributed species for reforestation. Previous studies have already associated the potential range of species with their resilience or adaptability to climate change (Booth 2016). However, while a selection based on species potential range size would simplify the selection and allow greater resilience, it would also lead to greater homogenisation of restored landscapes (Wang et al. 2019). This, in turn, would also decrease the potential for biodiversity with negative consequences that are difficult to foresee. Given the tendency of co-occurrence among tree species, one possible possibility could be to restore a mix of tree species with at least one broadly ranged species and a set of co-occurring small-range species. Heterogeneous structure from a broader diversity of trees can increase structural complexity and favour biodiversity (Díaz-García et al. 2017). Mixing broader and more limited ranged species might potentially be an option for promoting ecological resilience of the community even under climate change. However, the modelling applied in this study is also associated with uncertainties and the use of habitat suitability maps should be done jointly with expert based local knowledge (Reside et al. 2019) to improve the quality of the management decision.

Being able to map current and future distributions of a large number of species at the global scale requires making limiting assumptions and coping with uncertainty. The application of species distribution models under climate change is associated with cumulated levels of uncertainty (Brun, Thuiller, et al. 2020), especially since important mechanisms are not considered within correlative models (Case and Lawler 2017). Hence, model intercomparisons as well as comparisons with independent datasets are important for validation before applying them to management decisions (Barnard et al. 2019). Global databases used to map tree species distribution typically display important spatial bias (Beck et al. 2014) and our occurrence data in some regions lacked spatial accuracy. The number of regions considered for the case study is limited and may summarize the complexity of the different biomes somewhat simplistically. We estimated future climate based on downscaled projections of several state-of-the-art climate models used by the IPCC (CMIP6) but still these projections are associated with substantial spatial and temporal uncertainty (Yazdandoost et al. 2021) and do not account for extreme events such floods, fires, and droughts or floods which can cause large tree mortality (Brun, Psomas, et al. 2020). While it was possible to validate the map with other data for Europe, the evaluation of the quality of the projections for other regions is more difficult, making the quality of the maps in the corresponding regions more uncertain. This uncertainty should be appropriately communicated to the stakeholders for the final use of species habitat suitabilities (Zurell et al. 2020).

## Conclusion

There is no straightforward condition for planning ecosystem restoration strategies within a given region, with a wide range of practices having different implications for local biodiversity and human wellbeing (R. L. Chazdon and Guariguata 2018). For active restoration initiatives, the decision of which combinations of tree species to consider for any region requires consideration of the social context, the local ecological heterogeneity, but also how the current ecological conditions will shift under climate change. Here, we show how machine learning and high computing performance can support the mapping of thousands of tree species. We focused mainly on the case of restoration of ecosystems, but reforestation could benefit biodiversity and provide services beyond strict conditions of restorations (Bhagwat et al. 2008) and should consider a broader landscape context (Temperton et al. 2019) and service demands (Brancalion and Holl 2020). In our analyses, we excluded non-native tree range, but many trees that are producing benefits including fruits are planted beyond their native range, and planting those could also promote social and economic benefits to local communities (Cerullo and Edwards 2019). Moreover, not all tree species considered can accommodate agroforestry. Hence, for broader applications, there is a large potential of combining the species range maps as produced here with trait information from various databases (Kissling et al. 2019; Kattge et al. 2011), which together can support the management of natural or semi-natural ecosystems. In addition, while the projected suitability maps provide general indication to facilitate local decision making, species selection should take real consideration of local knowledge where possible for guiding responsable land management practices (Osorio-Salomón et al. 2021). Finally, the maps developed in our study can serve beyond restoration and support the conservation of threatened tree species globally (Fremout et al. 2020) and inform a variety of environmental programs.

## Acknowledgements

This project was partly financed by the Internal WSL grant “Treemap” to LP. DNK and NEZ acknowledge funding from the Internal WSL grant “ClimEx”, the BiodivERsA project (FeedBaCks) with the national funder Swiss National Science Foundation (20BD21_193907). DNK acknowledges funding from the BiodivERsA project “Futureweb” with the national funder Swiss National Science Foundation (20BD21_184131). NvT, JvdH and TWC acknowledge funding from DOB Ecology.

## Methods

We combined python and Google Earth Engine (Gorelick et al. 2017) to generate a pipeline for the high resolution mapping of species for informing restoration. Each species distribution is modelled separately. After data cleaning, the pipeline consisted of two parts: (1) constructing the geographic range in which the species is considered, and (2) modelling ecological suitability with an ensemble machine learning model, allowing us to map the species distribution under climate, as well as multiple climate projections.

### Data sources and cleaning

Tree species occurrence data was downloaded from 13 different online databases, published and unpublished datasets (Table 1). We then merged the occurrences of all databases and removed duplicates. We used the global tree database of GlobalTreeSearch (Beech et al. 2017) which consists of 60,111 tree species to filter relevant species to only consider tree species for our analysis. There were between 3 and 895,762 observations per species, with a median number of observations per species of 837, making up a total of almost 20 million (19,969,481) observations for 4,172 species. The observations were aggregated to the 30-arc second pixel level to match the resolution of the predictor layers. Observations that fell off the pixel-grid used for the predictors were removed (e.g., observations close to a coast that are aggregated to a pixel in the water).

### Range

For each species, we constructed a geographic range that was used to select observations to be included in the model training, thus excluding observations of non-native invasive species. The same range was used for model projection to prevent predictions outside the native range. The range considered for each species was constructed based on the reported native range in the GlobalTreeSearch database (Beech et al. 2017) and the location of the observations.

The reported native range consists of the geometries of the countries in which the species is native, with a 1000 km buffer to compensate for potential gaps in the GlobalTreeSearch database and to allow for the species to spread under projected future climates. Particularly when the reported native range contained large countries, such as Russia, Canada or Brazil, it was important to exclude the areas in which no observations were found. For this purpose, we intersected the reported native range with a range constructed around the ecoregions in which there were observations that had at least 3 other observations within 1000 km. For small ecoregions (bounding box less than 1000 km in width or length), the ecoregion with a 1000 km buffer was included in the range. For larger ecoregions, a 1000 km buffer around the part of the ecoregion that is within 200 km of an observation was included in the range.

For computational purposes, if there were more than 10,000 observations, these were spatially aggregated for the construction of the range.

### Training data

The training data consisted of the observations that fall within the considered range, marked as presence = 1, and pseudoabsences, randomly sampled within the considered range, marked as presence = 0. The number of points in the training data was capped at 20,000 to assure the model could run in Google Earth Engine without running out of memory. When necessary, the observations were randomly sampled from the full set. The number of sampled observations and pseudoabsences in the training data (n_obs, training_ and n_PA,training_) depended on the number of observations (n_obs_) (Table 2).

**Table 2.** Number of observations and pseudoabsences included in the training data (n_obs, training_ and n_PA,training_) dependent on the number of observations (n_obs_).

| $n_{\text{obs}}$ | $n_{\text{obs,training}}$ | $n_{\text{PA,training}}$ |
| --- | --- | --- |
| $n_{\text{obs}} \geq 10,000$ | 10,000 | 10,000 |
| $1,818 \leq n_{\text{obs}} < 10,000$ | $n_{\text{obs}}$ | $20,000 - n_{\text{obs}}$ |
| $500 \leq n_{\text{obs}} < 1,818$ | $n_{\text{obs}}$ | $n_{\text{obs}} * 10$ |
| $n_{\text{obs}} < 500$ | $n_{\text{obs}}$ | 5,000 |

### Environmental variables and climate change scenarios

For species distribution modelling, we used nine environmental variables related to climate and soil conditions as predictive variables. Climate variables, including average annual temperature, temperature seasonality, annual precipitation, precipitation seasonality, growing season length and net primary production, were obtained from CHELSA V2.1 (Karger et al. 2017). These factors represent basic resource requirements, metabolic modifiers or disturbance constraints to plant growth and survival. Soil variables, including pH, coarse fragment content and silt content, were obtained from SoilGrids (Hengl et al. 2017). We extracted all variables at a 30 arc-second resolution and projected them to the World Geodetic System 1984 (EPSG 4326) projection.

### Species distribution modelling

For each species, we fitted an ensemble model consisting of two random forest (RF) and two generalised boosted models (GBM), each with two different complexity levels with regard to model formulation (Brun, Thuiller, et al. 2020). The final distribution map was then achieved by taking the average over the predictions of the four individual models. We only modelled species for which at least 20 observations were available, leading to 3,987 models in total. To avoid overfitting when modelling rare species, we limited the number of predictor variables used in each model based on the number of observations for the corresponding species. We ensured that the number of observations available was at least 10 times the number of predictors used. For species that had less than 90 observations, we selected the floor(n_obs_ / 10) most important predictors for that species by training the ensemble model with all predictors and computing variable importance. Otherwise, the 9 predictors were used for modelling.

The model output was probabilistic, and we used a 3-fold cross-validation with random fold assignment to determine the optimal threshold to transform the probabilistic output to binary, by assessing the threshold maximizing the true skill statistic (TSS).

Finally, the ensemble model was trained on the full training set and predictions were made on global maps of the covariates. While soil variables were kept constant, we performed the predictions on ten sets of climate variables, one of current climate (1981-2010), and nine of predicted future climates. We used the mean climate conditions over three time periods (2011-2040, 2041-2070 and 2071-2100) and three different socio-economic pathways (ssp’s) representing a full climate change mitigation scenario (SSP 1.26), a middle of the road scenario (SSP 3.70), and a worst case scenario (SSP 5.85). We used five different global circulation models (GCMs) from the sixth coupled model intercomparison project (CMIP6) that where bias-corrected (Lange 2019) for the intersectoral-impact model intercomparison project (ISIMIP) (Frieler et al. 2015)

